# Strategic *in vitro* bioaugmentation of indigenous *Actinobacteria* isolated from east Kolkata wetlands (Ramsar site) to degrade highly toxic waste engine oil in free and emulsified state

**DOI:** 10.1101/2022.08.14.503898

**Authors:** Souptik Bhattacharya, Sirsha Putatunda, Ankita Mazumder, Dwaipayan Sen, Chiranjib Bhattacharjee

## Abstract

The present study investigates the bioremediation of waste engine oil at both dissolved or being suspended in oily wastewater using an actinobacterium, *Gordonia terrae* DSM 43249 strain isolated from East Kolkata wetlands. The isolated strain was found capable of sustaining in highly toxic oil contaminated wastewater and simultaneously can efficiently biodegrade the pollutants in both simulated fresh and marine water system at optimized environmental conditions. Moreover, in order to understand the effect of physical presence of oil in oily wastewater on the bioremediation process, three types of simulated oil-water forms were studied: water with free waste engine oil, oil-water mixture in the form of coarse emulsion and microemulsion. It was observed that the percentage degradation became maximum with the microemulsion form (72.73%) followed by the coarse one (65.45%). The minimum percentage degradation of 39.74% was seen with the free oil. Statistical interpretation also revalidates the experimental observations, showing that the oil percentage degradation is much sensitive to the oil and water composition in an oily-water system (F=772.64> Fcritical =5.143). Hence, it is presumed from the present study that such a high percentage degradation of oil especially, when oil is thoroughly mixed with water, can be considered as one of the potential applications for oil treatment such as during oil spillage using *G*.*terrae* DSM 43249.

**Graphical Abstract:** 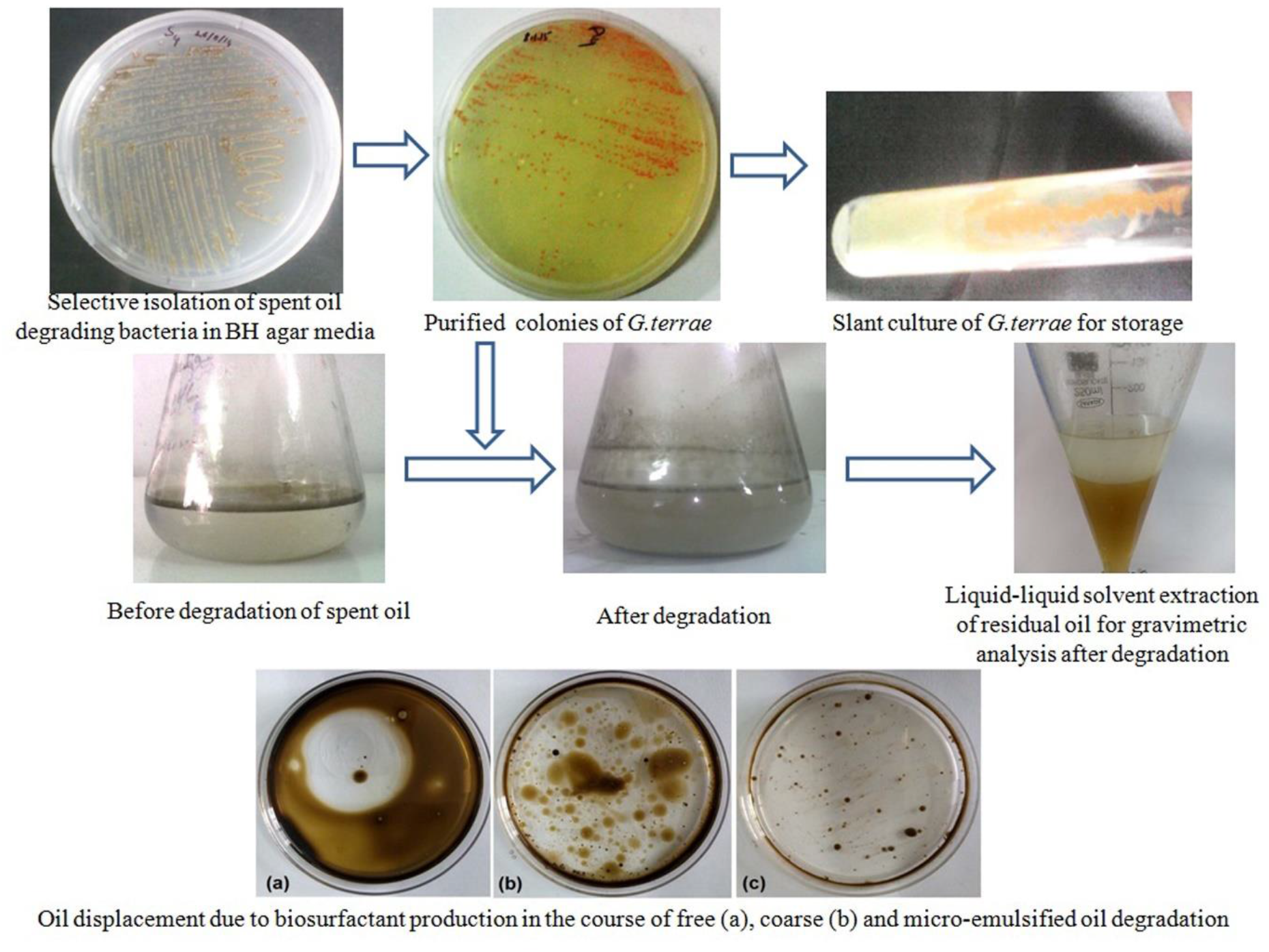

## 1. Introduction

Widespread urbanization in recent days increases the volume of wastewater generation either in the form of industrial effluent or wastewater from household activities [1]. Automobile sectors have been considered as one of the most common contributors to oily wastewater pollutants. The predominant components of these oily effluent are primarily waste engine oil, greases, lube oils, detergents etc. Among them, oil and grease are the most common and key contaminants imposing a serious threat to the environment [2]. Waste engine oil is heavier and sticky compared to new blended oil, and contains varied toxic pollutants like polychlorinated biphenyls (PCB), polyaromatic hydrocarbon (PAH), heavy metals, etc. Although, a small fraction of this waste engine oil is further reused, while major portion is disposed of as effluent without any proper treatment, which is a major challenge worldwide [3]. These pollutants can accumulate and persist in the environment for a long duration of time without any further degradation [4]. Even with trace waste engine oil in water body, it can adversely affect the aquatic ecosystem due to bioaccumulation. Thus, when the higher hierarchy organism of food chain such as human once consumes these aquatic species, high level of hazardous compounds gets transferred through bioaccumulation and impose their adverse health impacts.

Hence, a proper treatment of these contaminated oily waste streams is highly recommended. However, the main intricacy associated with the treatment of oil contaminated waste stream is the form of oil in which it exists in bi-phasic oil-water mixture. Oil can persist in wastewater mainly in two forms, free oil and emulsified oil. Moreover, depending on the size of dispersed phase, the stability of emulsion differs. There are two types of emulsion – one, kinetically stable coarse emulsion having size of 0.05-100 µm for the dispersed phase and another is thermodynamically stable, microemulsion, having reduced size droplets of radii 1-100 nm [5]. Several conventional treatment processes were found efficient enough to remove free oil, but they cannot ensure their efficiency to oil-water emulsion separation [6]. In this context, bioremediation technology was envisaged as an efficient, economic and eco-friendly strategy to enhance the removal of hydrocarbons from the waste stream [7]. The added advantage of this technique over conventional ones is that it converts the pollutants into less toxic components and often completely degrades the pollutants into their natural elemental forms. It was studied from several research outcomes that bioremediation involves low recurring investment, which in turn reduces the treatment cost per unit volume of wastewater generation [8,9]. The primary objective of this present study was to understand the efficiency of wetland microbiota’s bioremediation capability on the removal of oil from oil-water homogeneous mixture in control to free oil.

## 2. Materials and Methods

### 2.1 Reagent and standards

Purchases were made from Sigma Aldrich and Merck Chemicals Ltd for all the chemicals utilised in this investigation. Waste engine oil was gathered from nearby automobile repair shops. In accordance with the generally accepted bio-safety regulations, each experiment was carried out in triplicate.

### 2.2 Collection of wetland water for bacterial isolation

For the purpose of bacterial strain isolation, wetland water from ponds close to the East Calcutta Wetlands (Coordinate: 22.5135° N/88.4019° E) was collected. The Ramsar Convention recognised the East Kolkata Wetlands as a “wetland of international significance” on August 19, 2002 [10].

### 2.3 Isolation and identification of bacteria

The samples were inoculated in Bushnell-Haas (BH) broth media with the filter-sterilized waste engine oil as sole carbon source [11]. The isolation was done using the method proposed by H.M.M Ibrahim 2016 [12]. Selected pure isolate was then stored at 4°C for further study on oil degradation. A few phenotypic characterizations and the colony morphology study of the pure strain were carried out. Further, the strain was identified using 16s rRNA gene identification process from MTCC Chandigarh, India. The following sets of universal primers, 27F (5′-AGAGTTTGATCMTGGCTCAG-3′) and 1492R (5′-TACGGYTACCTTGTTACGACTT-3′), were used for polymerase chain reaction (PCR) amplification. Using a BLAST search on the National Centre for Biotechnology Information (NCBI) service, the 16S rRNA gene sequence alignment similarities were determined.

### 2.4 Evaluation of optimized growth condition for purified bacterial strain

The growth parameters (pH and temperature) of the isolated pure strain were optimized after studying the growth condition at varying range of pH from 5 to 10 and temperature from 25°C to 40°C. The sustainability of the isolated strain in fresh and marine environment was studied by conducting its growth in synthetic system resembling both the environment respectively. Ionic effect on the growth was assessed with the BH media mixed with three different salt systems (Na^+^ (55%) with Mg^2+^ (4%)), Ca^2+^ (1%) with K^+^ (1%) and sea salt). The estimation of bacterial growth was carried out by measuring the optical density (OD) at 600 nm.

### 2.5 Preparation and characterization of coarse- and micro-oil/water emulsion

Oil-in-water (o/w) coarse emulsion was prepared through mixing waste engine oil with DI water in the presence of SDS as surfactant in the ratio of 0.5:98.5:1 at a stirring speed of 100 rpm. Microemulsified wastewater was prepared by waste engine oil 25% (w/w), surfactant (Tween 80) 60% (w/w), co-solvent (n-butanol) in water 15% (w/w) under mild stirring at 25°C (fig. 1(a) and (b)). Droplet size and zeta potential were calculated using Malvern Zetasizer Nano ZS-90 (Malvern Instruments Ltd, UK). Further, the stability of both the coarse emulsion and microemulsion system were observed for 30 days at 35°C.

**Fig 1:**
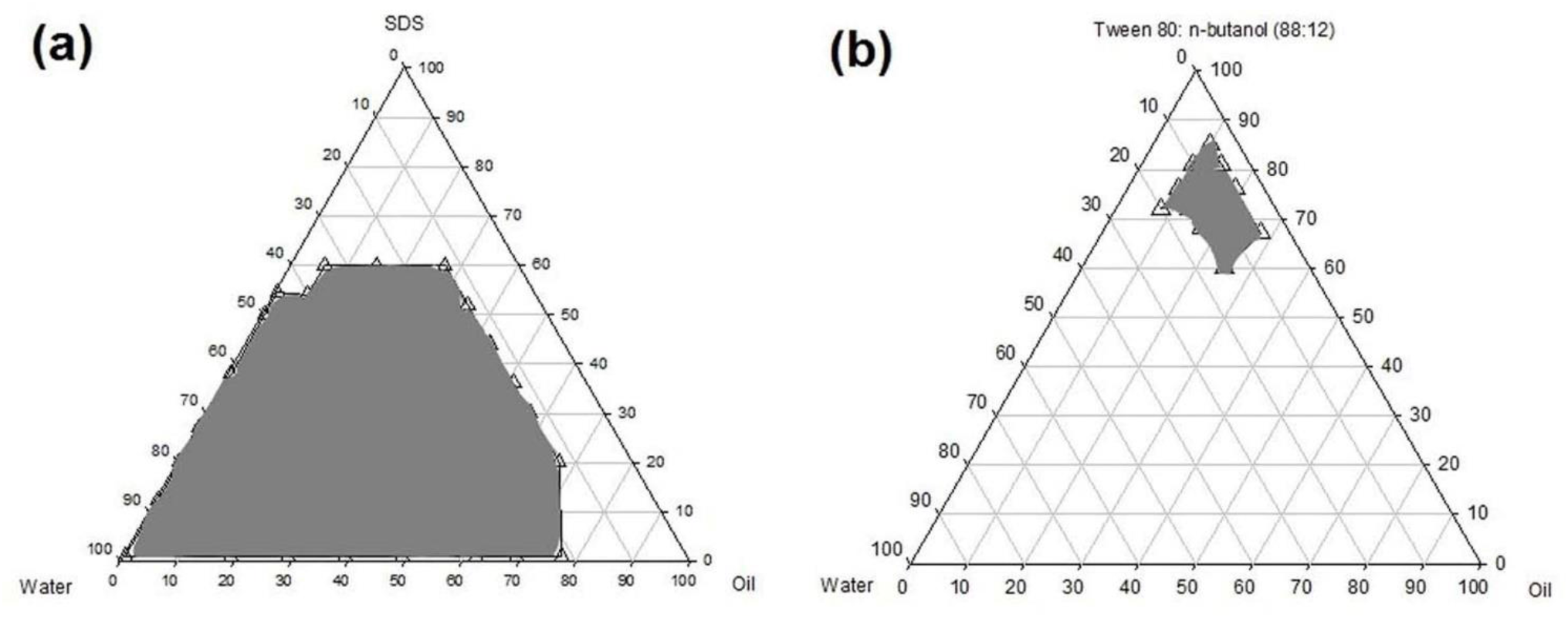
Ternary phase diagram where the portion within the marked boundary represent the positive formulation of (a) o/w coarse and (b) micro-emulsion

### 2.6 Biodegradation of free oil, emulsion and microemulsion using isolated strain

2% (v/v) of waste engine oil (either free oil form or oil in coarse emulsion or oil in microemulsion) and 1% (v/v) of pre-cultured overnight bacterial inoculums were mixed with 50 ml of BH broth media. The mixture was then incubated at optimum temperature and pH for 30 days at 120 rpm. After 30 days, the residual oil present in the system was extracted using 0.8 ml of CCl_4_ per ml of broth. Solvent was then completely evaporated using a vacuum oven. The gravimetric analysis was carried out to evaluate the percentage of consumed oil during the bacterial growth for each o/w system (Eq. 1).

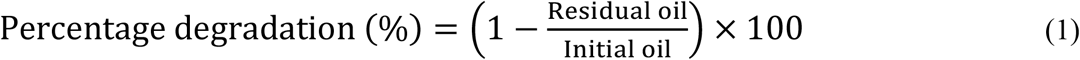

Surface tension of the BH media with three different oil systems were measured during the growth phase using stalagmometer (make: Borosil). Oil displacement study was carried out to confirm the presence of biosurfactant in the system [13]. Three Petri dishes, each containing 20 ml of BH culture broth were taken where 1 ml of free oil, 1 ml of coarse emulsion and 1 ml of microemulsion were added at the centre of each respective petridish. The diameter recorded of the clearing zone indicated the oil displacement activity of the biosurfactant produced.

## 3. Results and Discussion

### 3.1 Analysis of oil degrading microorganism

The strain was identified by 16s rRNA gene sequencing. The most probable closest match was found with *Gordonia terrae* DSM 43249, an actinomycete (Acc no. NR_118598.1), where the pair wise similarity was found 99.13% from NCBI MegaBlast search. A phylogenetic tree was constructed using iTOL V6 for the isolated strain (fig. 2). Moreover, phenotypic and morphological characteristics of the strain were shown in table 2.

**Fig 2:**
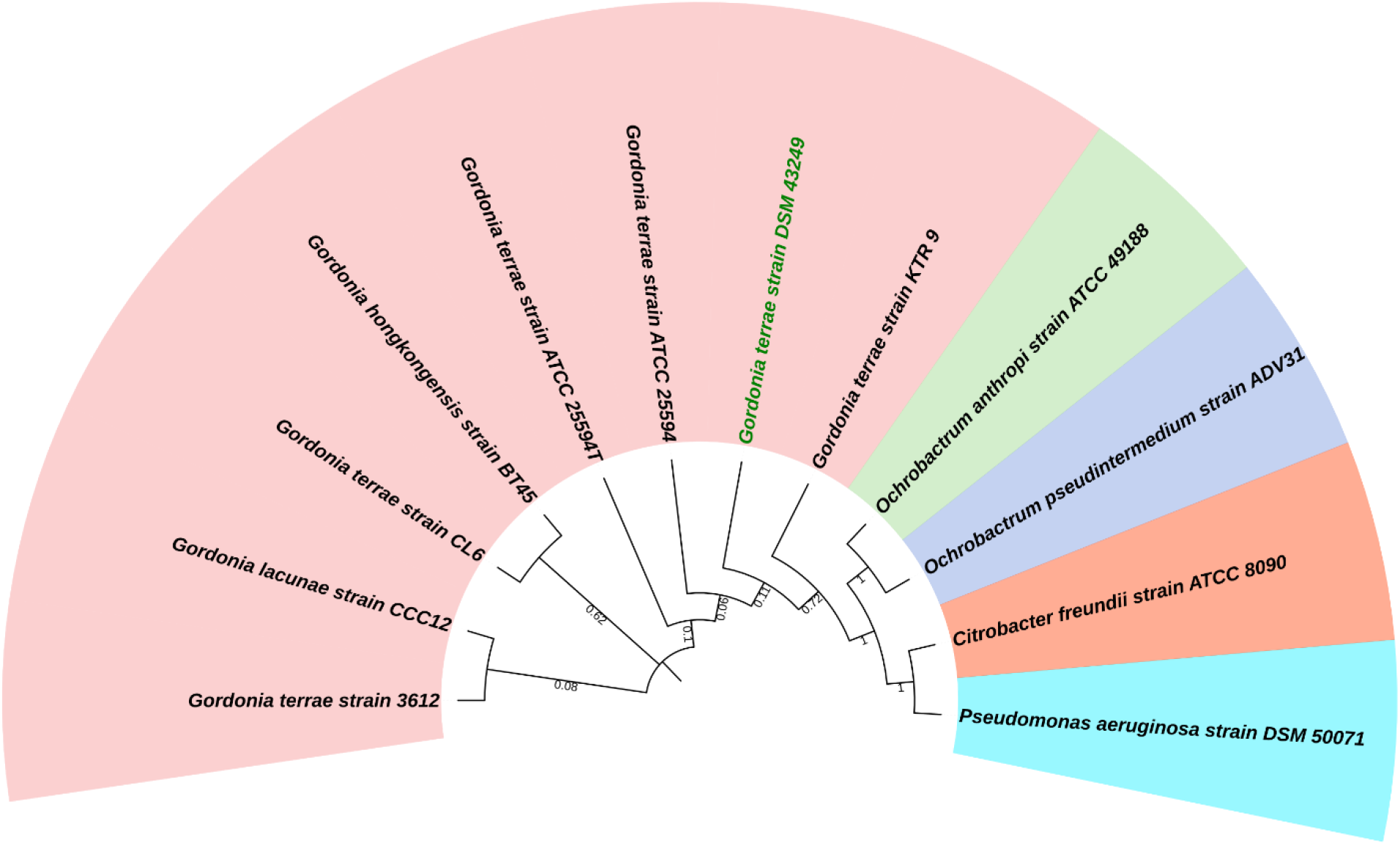
Phylogenetic tree of *G. terrae*

**Table 1:** Physicochemical characterization of oily wastewater

| Sl no. | Parameters | Sample site 1 | Sample Site 2 |
| --- | --- | --- | --- |
| 1 | pH | 5.3 | 5.7 |
| 2 | Total Suspended Solids (TSS)<br>(ppm) | 3793 | 2933 |
| 3 | Total Dissolved Solids (TDS)<br>(ppm) | 2770 | 2900 |
| 4 | Conductivity (mS/cm) | 0.78 | 0.92 |
| 5 | Salinity (ppt) | 0.398 | 0.472 |
| 6 | Total microbial load (cfu/ml) | $4.5 \times 10^7$ | $3.1 \times 10^7$ |
| 7 | Total oil degrading microbial<br>load (cfu/ml) | $3.1 \times 10^5$ | $3.0 \times 10^5$ |
| 8 | Oil & grease (ppm) | 56 | 74 |

**Table 2:** Phenotypic and morphological test of the isolated strain *G. terrae*

| Phenotypic Morphological Tests of *G. terrae* |  |  |  |  |  |
| --- | --- | --- | --- | --- | --- |
| Biochemical Tests | Gram's staining | Catalase test | Starch hydrolysis test | Motility test | Indole test |
|  | Positive | negative | negative | negative | negative |
| Colony Morphology Tests | Color | Type | Form | Elevation | Texture |
|  | Pink- Orange | Opaque | Circular | Pulvinate | Buttery |

### 3.2 FTIR analysis of the waste engine oil

Fig. 3 shows the FTIR spectrogram of the waste engine oil. Strong absorption band in between 2800 cm^-1^ and 3200 cm^-1^ is because of the aliphatic –CH vibration. Low intensity peaks in the range of 1600 cm^-1^ and 1800 cm^-1^ is because of the attached oxygenated functional group such as carboxyl and nitro group. The intense peaks in the range 1350 cm^-1^ to 1450 cm^-1^ shows the deformation for –CH_3_ and –CH_2_ bonds. The low intense peak at around 1150 cm^-1^ shows the peak for S=O group formation because of the sulphur present in the lubricating oil [14]. However, most of the sulphur has been burnt and emitted as SO_x_. Peak at around 700 cm^-1^ to 900 cm^-1^ mainly attributes to the peaks because of S-OR esters, P-OR esters. Peaks around 450 cm^-1^ to 550 cm^-1^ show the presence of iron oxide. After biodegradation the new peaks at 1736 cm^-1^ and 1701 cm^-1^ were found manifesting the presence of esters and carboxylic acids as major degraded products [15].

**Fig 3:**
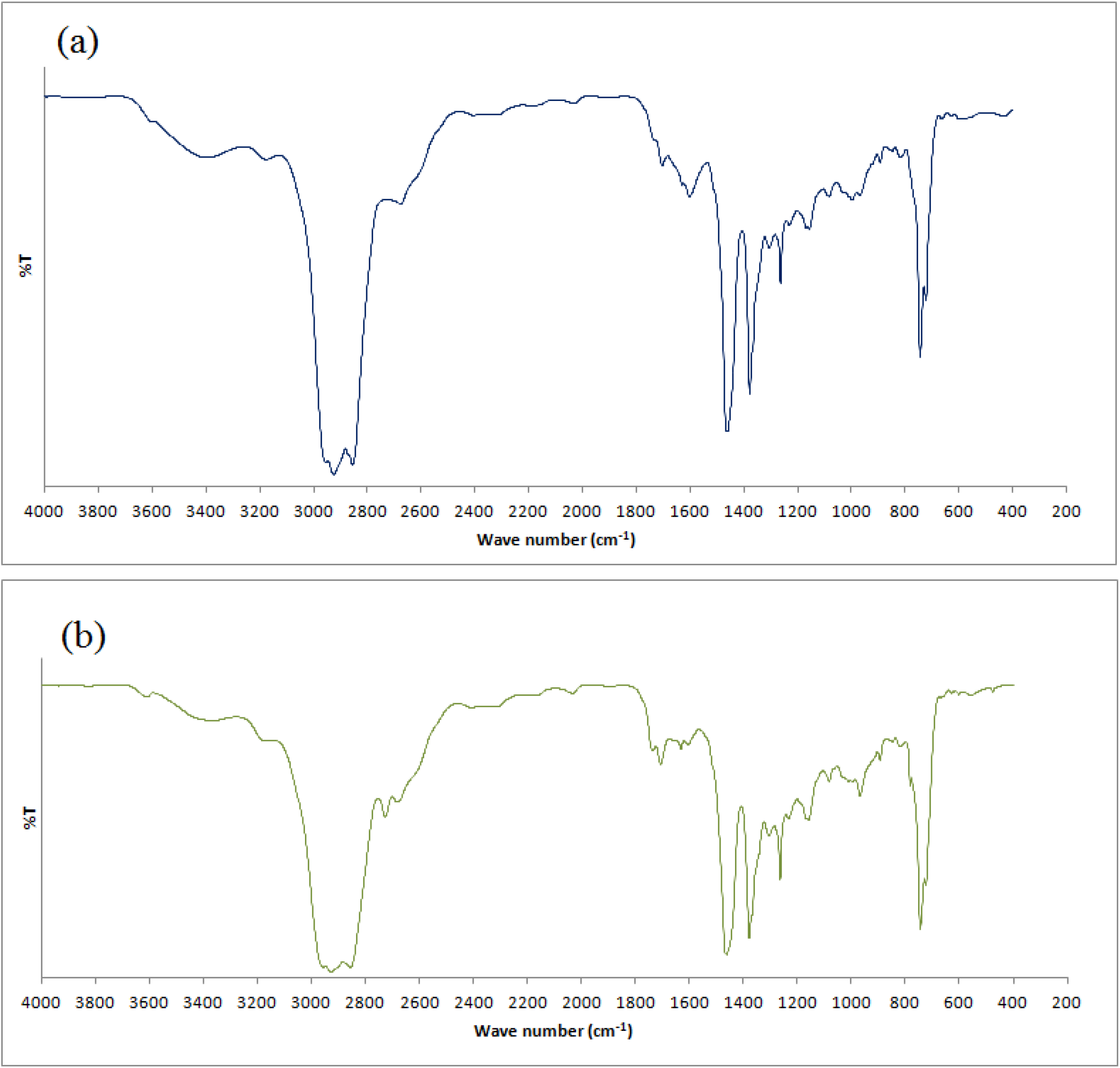
FTIR spectrogram of spent oil collected from automobile service shop. (a) Before biodegradation, (b) after biodegradation.

### 3.3 Zeta analysis for stability study of both coarse and microemulsion

The coarse emulsion droplets’ sizes were found large in the range of 500-800 nm, while for microemulsion, it was below 100 nm. Moreover, the zeta potential for both coarse and microemulsion were determined in the range of -50 to -80 mV, manifesting the abundance of negatively charged droplets [16]. In terms of stability, the coarse emulsion showed around 7-8% phase separation (fig. 4), whereas microemulsion didn’t show any separation attributing to its high stability [17].

**Fig 4:**
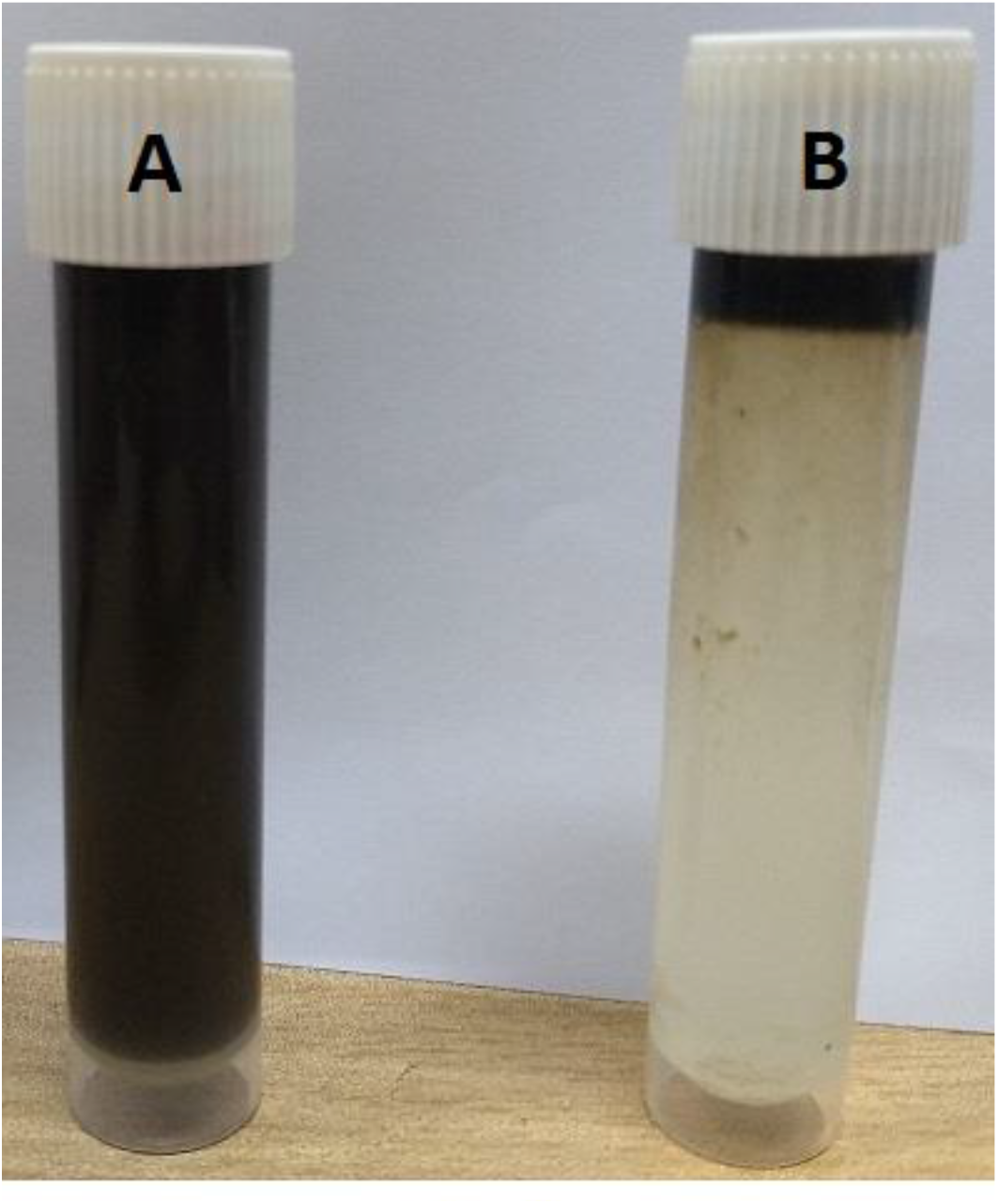
Stability test of coarse emulsion; (A) the emulsion before stability test and (B) the phase separated emulsion after 30 days of stability test.

### 3.4 Effect of temperature and pH for the bacterial growth

The bacterial growth was found maximum at pH 8 and 35°C (fig. 5) manifesting its mesophilic nature. One of the acclaimed observations is that the growth substantiates at a pH complies with the marine pH, which may be an additional benefit in remediation of oil from marine water.

**Fig 5:**
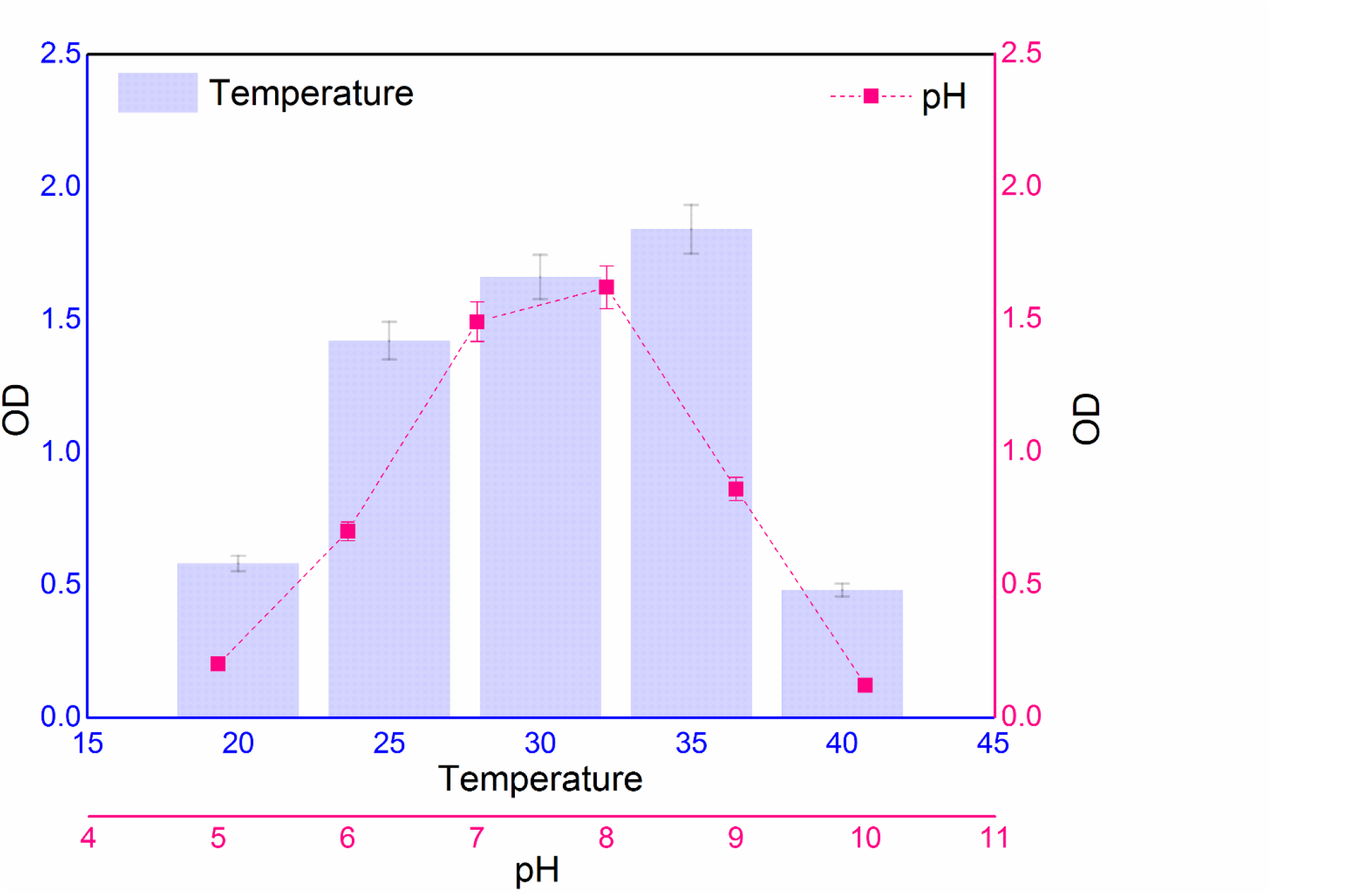
Effect of temperature and pH on the growth of *G. terrae*.

### 3.5 Effect of fresh and marine water environment on the bacterial growth

Isolated strain has portrayed its ability of sustaining in both simulated fresh water and marine environment. From fig. 6, it is evident that the strain had grown efficiently in fresh as well as in marine water with different ionic concentrations discussed previously. However, the growth of the strain is around 34.12% less in marine environment compared to fresh water medium. As *G. Terrae* was isolated from fresh water, it could not comply with the marine bacterial salt tolerant mechanism to maintain the same growth rate as in fresh water. Therefore, though the application of this strain for oily wastewater remediation will be mostly suitable for fresh water systems, it can also be used in case of marine oil spill.

**Fig 6:**
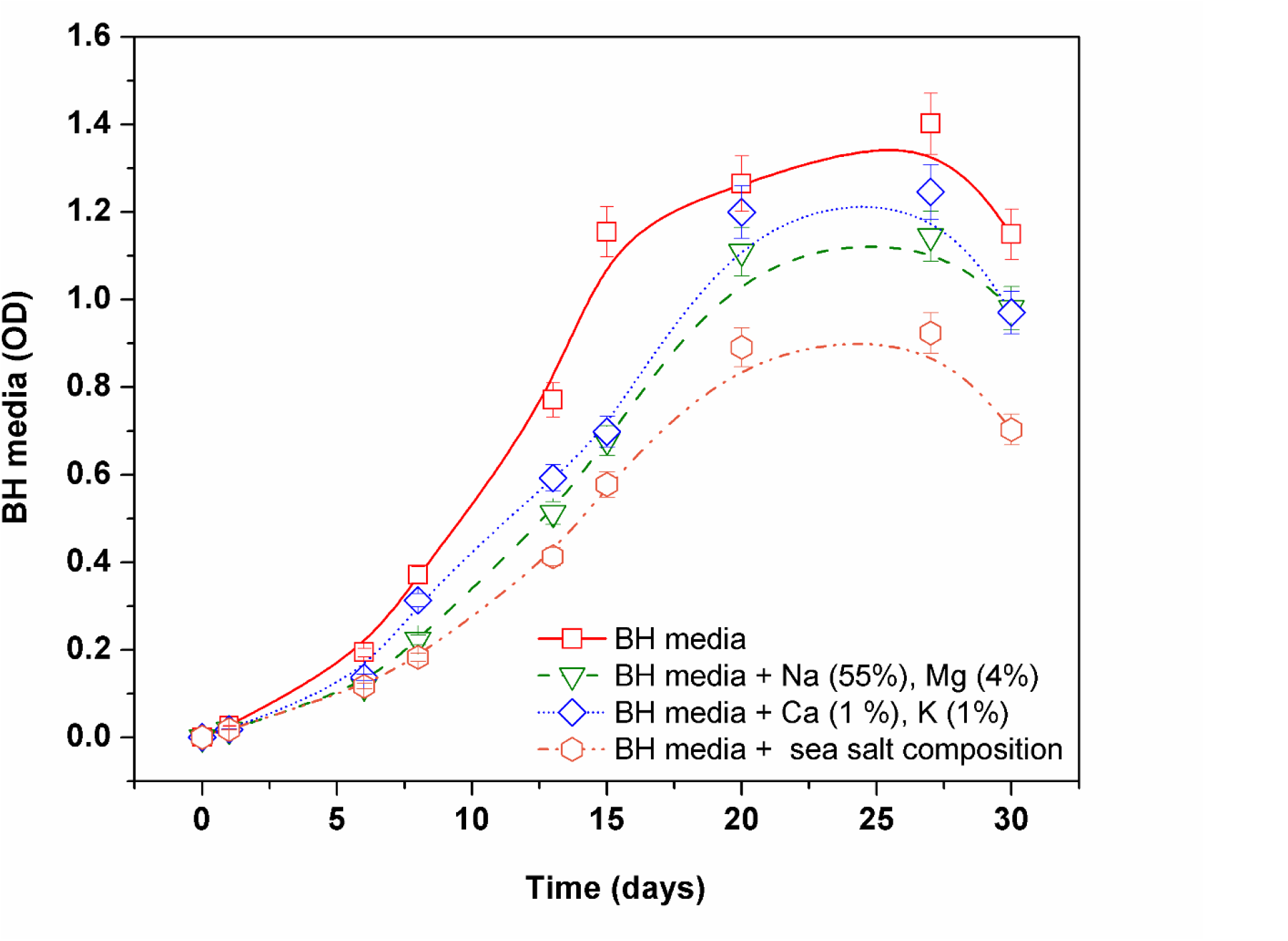
Growth study of *G*.*terrae* in different environmental aqueous systems.

### 3.6 Study on the degradation of free and emulsified oil using isolated strain

It is evident from fig. 7 that for both free and emulsions, the maximum biodegradation had taken place during the exponential growth phase of the bacteria. It was seen that bacteria degrades various hydrocarbons in preferential way depending on their structure. In general, the degradation follows as linear long chain alkanes (C_10_–C_40_)> branched chain alkanes> monoaromatics> cycloalkanes> PAHs [18]. Initially during the degradation process, the saturated fraction (e.g. n-alkanes) of the waste engine oil degrades, which yields an increased fraction of unsaturated hydrocarbons [19]. It was seen that that the degradation was maximum with the microemulsified oil (72.73±0.52%) and minimum with the free oil (39.74±0.25%). While, with the coarse emulsion the percentage degradation was 65.45±0.32%. Due to reduction in the size of the droplet the surface area per unit volume of the droplet increases facilitating more surface anchor with the bacteria. In the absence of nitrogen bacteria produce biosurfactants, a secondary metabolite to facilitate nutrient transport into the cell [20]. Such production of biosurfactants reduces the surface tension of the medium increasing the bioavailability of hydrophobic hydrocarbons to the microorganisms [13]. Fig. 8 shows the changes in surface tension of the medium with number of days for which growth was studied. From the figure, it is observed that the surface tension of both free and emulsified oil-water systems gets reduced substantially in the exponential growth phase of the strain (in between 6 to 18 days) due to maximum consumption of hydrocarbons along with the production of biosurfactants. Maximum production of biosurfactant was ascertained from relative standard deviations (RSDs) in the decrease of surface tension at exponential growth phase for all free, coarse emulsion and micro-emulsion. It was noticed that the minimum deviation was with the free (16%) and maximum with the micro-emulsion (31%) inferring increased bioavailability of oil fraction in micro-emulsion. Fig. 9 shows the disappearance of oil in all the three systems due to mixing because of biosurfactant production. The maximum disappearance of oil was achieved with the microemulsion broth due to elevated production of biosurfactant that ultimately increases the solubility of oil and enhances its uptake by the bacterium. Increased interfacial area in emulsion facilitates bioavailability of oil to microbes for consumption [21]. Moreover, the readily available surfactants present in the emulsion facilitates nutrient transport of water-insoluble substrates into the cell and thus manifests better biodegradation of emulsified oil systems [20]. Hence, in consequence, the degradation was maximum with the microemulsion compared to free oil. It shows around 1.8 fold more degradation compared to free oil. On the contrary, with coarse emulsion, the degradation percentage was 1.6 fold more compared to free oil. As the RSD between these two fold increases is only 7% attributing to the fact that metabolic activity of the strain on the emulsified system is not much different from each other.

**Fig 7:**
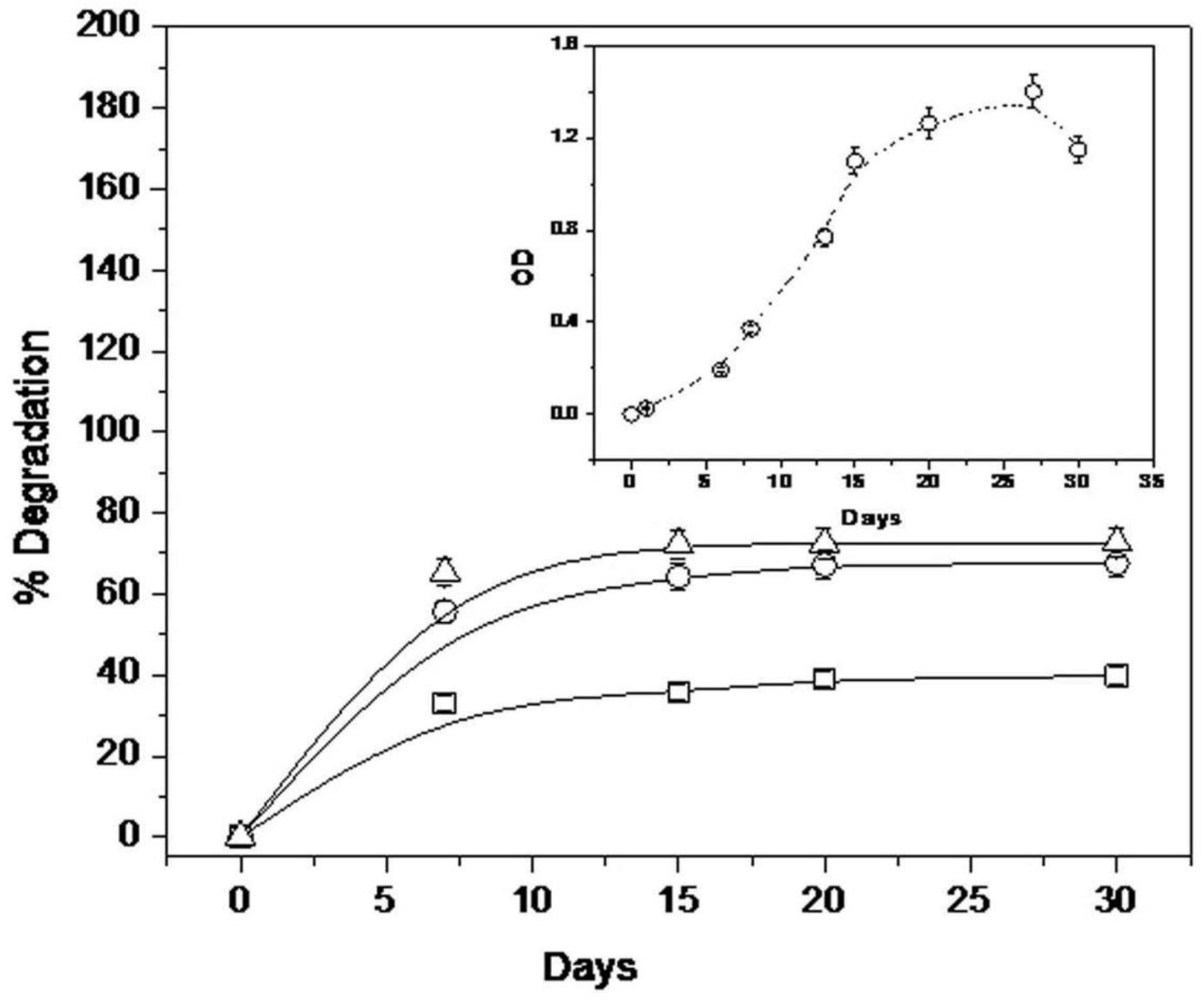
Variation of % Degradation oil with days for free oil (-□-), coarse emulsion (-o-) and microemulsion (-Δ -) (inset: Growth kinetics of *G. terrae*)

**Fig 8:**
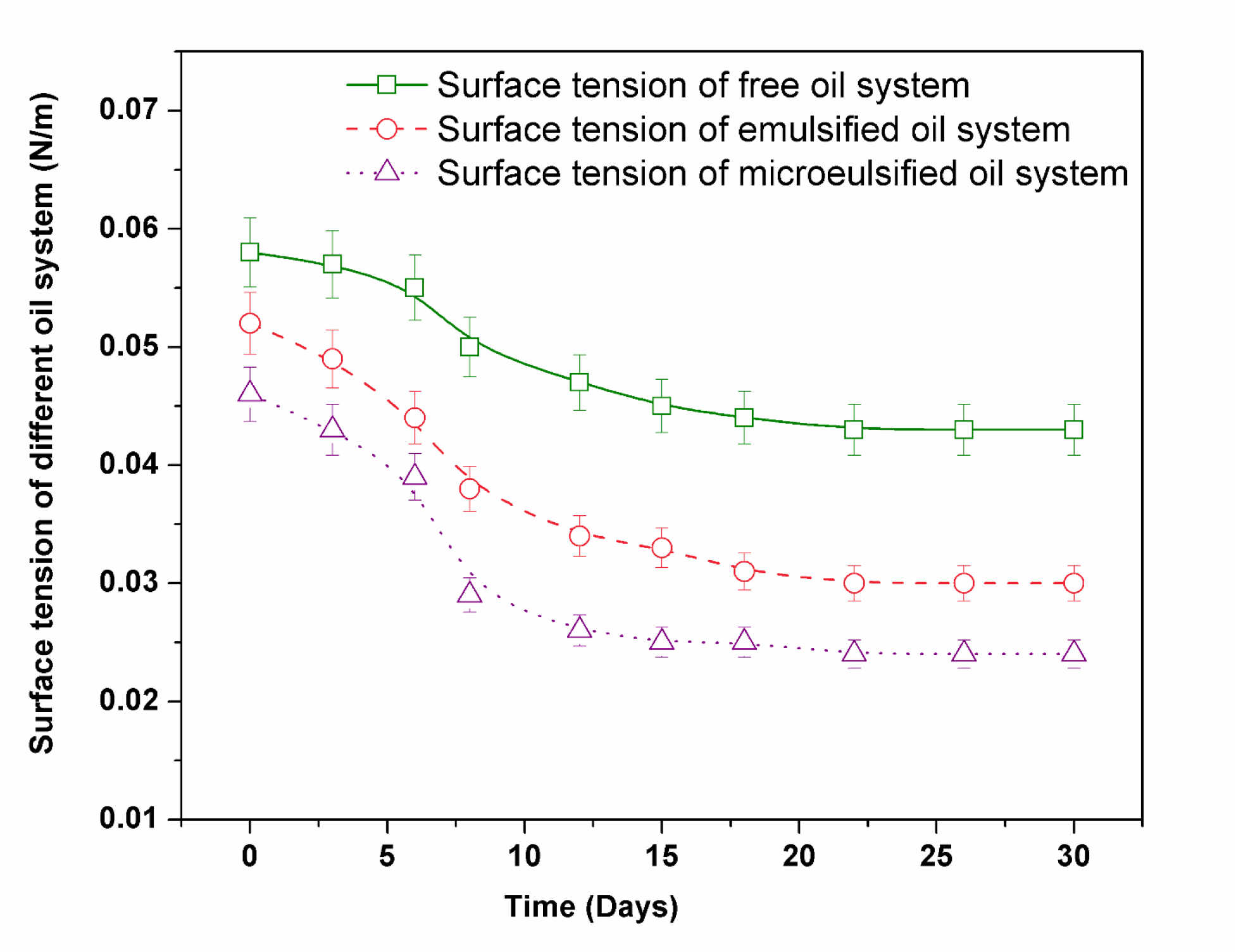
Variation of surface tension (N/m) in free, coarse and micro o/w emulsion

**Fig 9:**
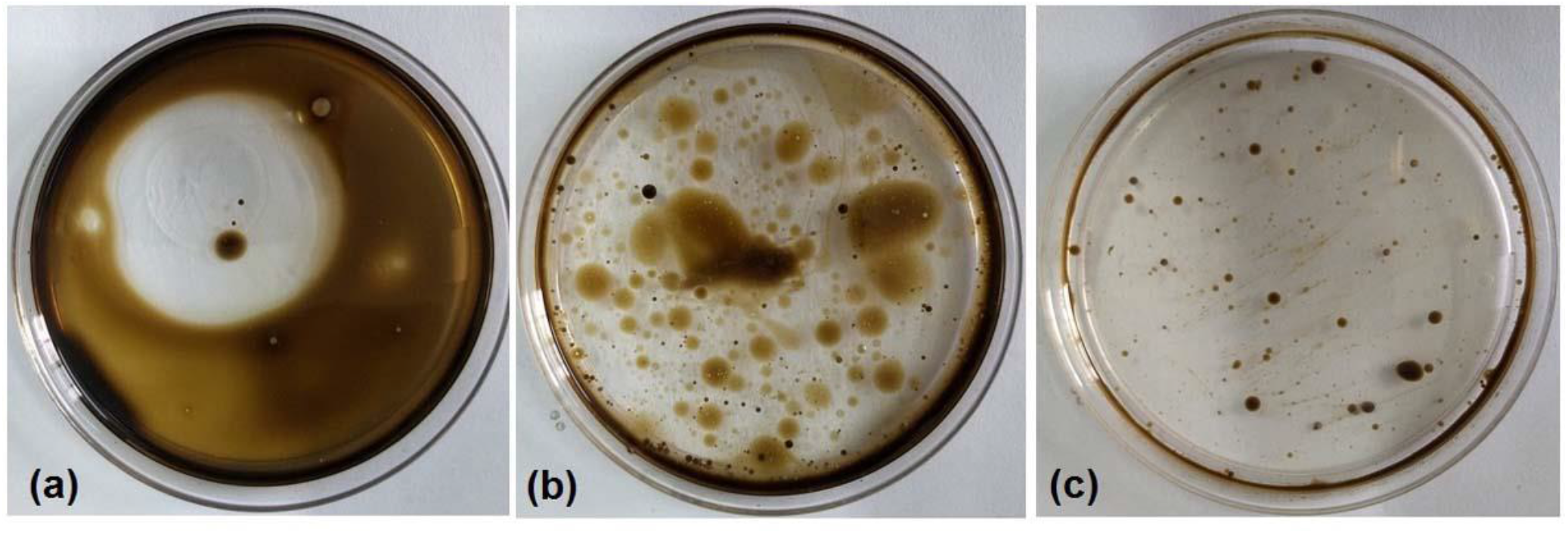
State of the substrate during bioremediation at the end of the growth phase: (a) free oil, (b) coarse, and (c) micro-emulsion.

### 3.7 Statistical significance test on the degradation percentage based on oil-water system using ANOVA

The one-way ANOVA test was carried considering three individual groups as “Oil”, “Emulsion” and “Microemulsion”. Table 3 illustrates the results of the ANOVA test, where the significance test was conducted with a 5% confidence interval. Table 3 shows that the F-value, defined as the variance in group means over mean of the variance within the group, is 772.65 (> F_Critical_=5.143253) between the groups. Such a large F value manifests that the degradation of oil depends on the nature of substrate, not on the metabolic activity of the strain *G. terrae* for a particular substrate. It is reassessed with the p-value (∼0<0.05) that the strain efficiency in bioremediation depends on the nature of the substrate, which is in this case either free oil or oil in emulsion.

**Table 3:** One way ANOVA test for determining the effect of the types of oily wastewater on the biodegradation efficiency

| *Source of Variation* | *SS* | *df* | *MS* | *F* | *P-value* | *F<sub>critical</sub>* |
| --- | --- | --- | --- | --- | --- | --- |
| Between Groups<sup>@</sup> | 1874.929 | 2 | 937.4644 | 772.6497 | 0.000000058 | 5.143253 |
| Within Groups<sup>#</sup> | 7.279867 | 6 | 1.213311 |  |  |  |
| Total | 1882.209 | 8 |  |  |  |  |

<sup>@</sup>: “Between groups” refers to different types of oil-water system (free oil, coarse and microemulsion)
<sup>#</sup>: “Within groups” refers to replications of analysis for a particular oil-water system

## 4. Conclusion

The present study emphasized on the bioremediation of waste engine oil, a major pollutant in oily wastewater using indigenous bacterial strain *Gordonia terrae*. It was seen that the strain’s metabolic activity during the degradation of oil from oily wastewater depends on the form of oil within water. The bacterial strain was much prompt in degrading waste engine oil with the emulsified oil compared to free oil present at the surface of the water. Especially the form of microemulsified oil shows a bit higher percentage degradation of oil with the strain. However, the difference in the percentage degradation of oil is not significantly different from each other in case with coarse and microemulsified oil. The coarse emulsified and microemulsified oil degradation was found to be 65% and 83% higher than free oil respectively. The increased surface area of waste engine oil during biodegradation of the emulsified systems facilitates higher degradation amount of emulsion than free oil system. Hence, bioremediation using indigenous Actinomycete *Gordonia terrae* was found to be a promising bio agent for treating emulsified oily stream comprised of free and emulsified waste engine oil phase.

## Acknowledgement

The authors acknowledge the senior research fellowship and contingency grant awarded to SB by CSIR, India (File No. 09/096(0879)/2017-EMR-I) which provided necessary support to carry out the investigation.

## Author Contribution Statement

**Souptik Bhattacharya:** Conceptualization, Investigation, Data curation, Formal analysis, Methodology, Software, Visualization, Funding acquisition, Writing – original draft, Writing – review & editing. **Sirsha Putatunda:** Conceptualization, Data curation, Funding acquisition, Supervision, Validation, Formal analysis. **Dwaipayan Sen:** Methodology, Software, Investigation, Visualization, Writing – review & editing. **Ankita Mazumder:** Investigation, Data curation, Formal analysis, Methodology. **Chiranjib Bhattacharjee:** Conceptualization, Funding acquisition, Project administration, Resources, Supervision, Validation, Visualization, Writing – review & editing.

